# Pax6 loss alters the morphological and electrophysiological development of mouse prethalamic neurons

**DOI:** 10.1101/2021.03.22.436455

**Authors:** Tian Tian, Idoia Quintana-Urzainqui, Zrinko Kozić, Thomas Pratt, David J. Price

## Abstract

Pax6 is a well-known regulator of early neuroepithelial progenitor development. Its constitutive loss has a particularly strong effect on the developing prethalamus, causing it to become extremely hypoplastic. To overcome this difficulty in studying the long-term consequences of Pax6 loss for prethalamic development, we used conditional mutagenesis to delete Pax6 at the onset of neurogenesis and studied the developmental potential of the mutant prethalamic neurons in vitro. We found that Pax6 loss affected their rates of neurite elongation, the location and length of their axon initial segments and their electrophysiological properties. Our results broaden our understanding of the long-term consequences of Pax6 deletion in the developing mouse forebrain, suggesting that it can have cell autonomous effects on the structural and functional development of some neurons.

**SUMMARY STATEMENT:** Pax6 impacts neurite extension, axon initial segment properties and ability to fire normal action potentials in maturing neurons, revealing actions extending beyond those previously characterized in progenitors.

## INTRODUCTION

From a very early stage, the neuroepithelium is patterned by the regional expression of transcription factors that specify the subsequent development of each region. The extent to which the actions of these transcription factors influence the later development of the functional properties of neurons is much less clear.

We studied the transcription factor Pax6, a well-known regulator of early neural development (Cvekl and Callaerts, 2017). Pax6 expression in the mouse neuroepithelium is first detected soon after neural tube closure and continues in specific forebrain progenitors, where it has pivotal functions in diverse early developmental processes (Hanson and Van Heyningen, 1995; Engelkamp *et al*., 1999; Mi *et al*., 2013; Manuel *et al*., 2015; Cvekl and Callaerts, 2017). In most of these regions, Pax6’s expression is lost in cells exiting the cell cycle during neurogenesis (Duan *et al*., 2013). Unusually, strong Pax6 expression is retained by many post-mitotic neurons in the prethalamus (Caballero *et al*., 2014). Constitutive deletion of Pax6 all but prevents the formation of the prethalamus, precluding an analysis of the consequences of its loss for developing prethalamic neurons (Stoykova *et al*., 1996; Warren and Price, 1997).

We re-examined our previously-reported RNAseq dataset of changes in prethalamic gene expression following acute Pax6 deletion at the onset of neurogenesis (Quintana-Urzainqui *et al*., 2018) and found upregulated expression of genes involved in neuronal morphogenesis and ion transport. To investigate the possibility that Pax6 loss might affect the ability of prethalamic neurons to acquire their normal structural and functional properties, we studied the developmental potential of prethalamic neurons with the same acute Pax6 deletion. We used dissociated culture, thereby increasing the likelihood of detecting cell autonomous effects. We found that the neurites of Pax6-deleted prethalamic neurons grew at abnormal speed, their axon initial segments (AISs) tended to be longer and to extend further from the soma and their electrophysiological properties were altered. Our results indicate that in addition to its role in early prethalamic progenitors, Pax6 is also required in the later structural and functional development of prethalamic neurons.

## RESULTS AND DISCUSSION

### Prethalamic Pax6 deletion caused upregulation of genes involved in neuronal morphogenesis and ion transport

At E13.5, Pax6 is expressed in cortical, thalamic and prethalamic progenitors and in a population of prethalamic neurons (Figure 1A-A’). We interrogated an existing RNAseq dataset showing significant transcriptional changes (adjusted p < 0.05) in E13.5 prethalamus after acute deletion of Pax6 from E11.5, which is after prethalamic neuroepithelial specification and around the time of onset of prethalamic neurogenesis (Quintana-Urzainqui *et al*., 2018). Gene ontology (GO) term enrichment analysis on these genes revealed 495 upregulated and 125 downregulated GO terms (Supplementary Table 1). Among the top 200 upregulated GO terms, 25 related to neuronal morphogenesis and 12 to ion transport. After removing child terms, 14 GO terms were related to neuronal morphogenesis and 7 were related to ion transport (Figure 1C-D, Supplementary Table1).

**Figure 1.**
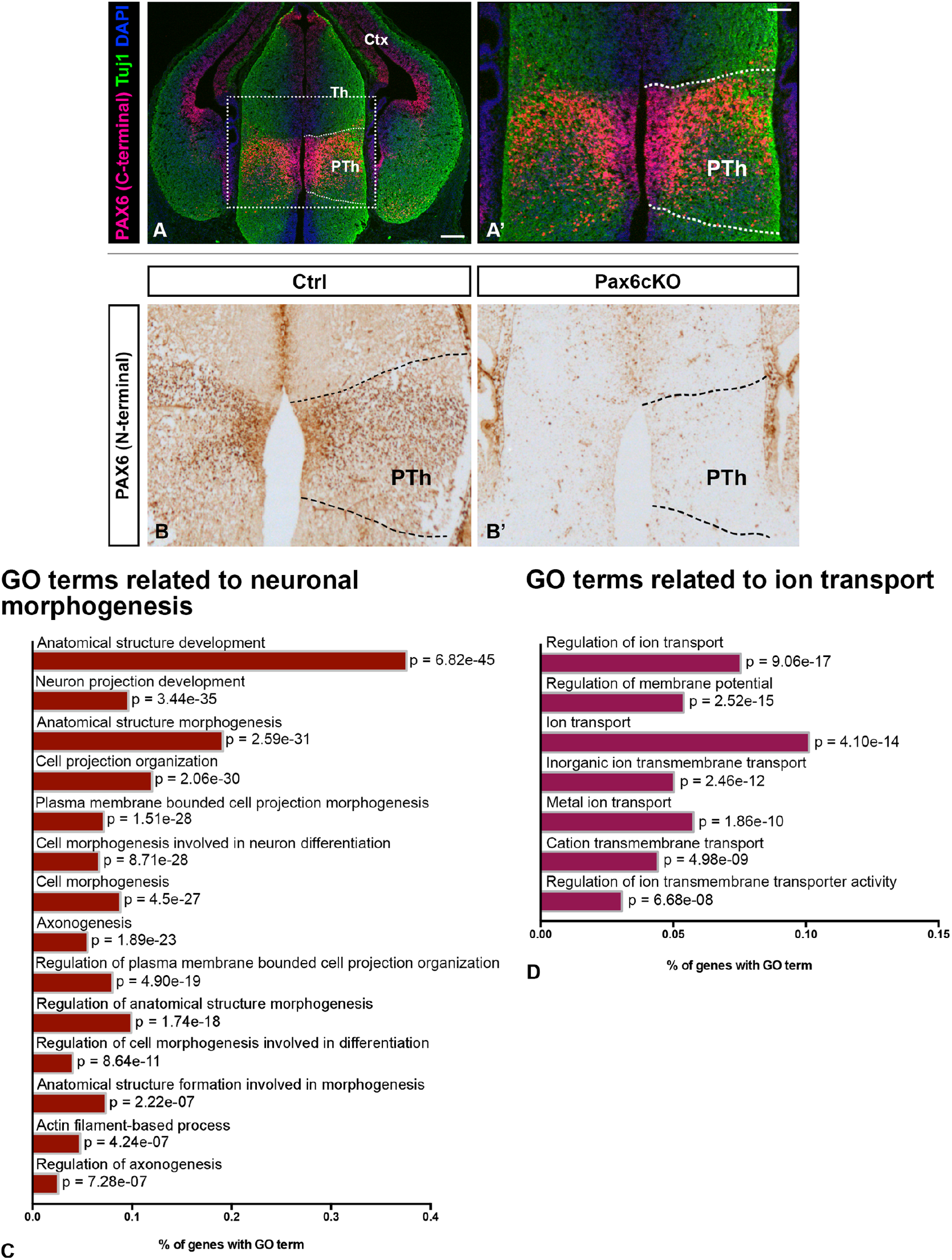
Pax6 deletion in the prethalamus caused upregulated expression of genes involved in neuronal morphogenesis and ion transport related GO terms. (A-A’) Immunohistochemistry showing Pax6 expression overlapping with Tuj1, a marker for postmitotic neurons, in the E13.5 prethalamus of the control littermate. PTh, prethalamus, Th, thalamus, Ctx, cortex. (B-B’) Pax6 immunohistochemistry at E13.5 showing CAG^CreER^-induced loss of Pax6 in the prethalamus following tamoxifen administration at E9.5. (C) The top 14 most highly enriched, non-redundant GO terms related to neuronal morphogenesis and (D) the top 7 most highly enriched, non-redundant GO terms related to ion transport in the Pax6 cKO prethalamus. Scale bar: (A) 250 μm; (A’, B-B’) 100μm. See also Supplementary Table 1.

Pax6 loss upregulated genes important for the actin and microtubule cytoskeleton during neuritogenesis, for axon specification and for neurite elongation (Supplementary Table 2). In other systems, upregulation of these effector genes increased actin nucleation, actin bundling, microtubule assembly and bidirectional intracellular transportation, which have been shown to augment axon/neurite elongation (Witte and Bradke, 2008; Stiess and Bradke, 2011; Flynn, 2013; Sainath and Gallo, 2015). Pax6 loss also upregulated genes involved in relaying extracellular signals to both the actin and microtubule cytoskeleton and genes encoding several components of the PAR3/6 complex, which plays a vital role during the establishment of neuronal polarity (Supplementary Table 2) (Nishimura *et al*., 2004; Shi *et al*., 2004; Barnes and Polleux, 2009; Lalli, 2014).

The AIS is located at the most proximal end of the axon and is the site where action potentials usually initiate (Kole and Stuart, 2008; Kole *et al*., 2008; Rasband, 2010; Leterrier, 2018). Genes encoding the cytoskeletal components of the AIS were also upregulated (Supplementary Table 2), including the cytoskeletal scaffold protein AnkyrinG (AnkG, also known as Ank3) and βIV Spectrin (Sptbn4). In neurons, AnkG is restricted to the AIS and nodes of Ranvier, where it tethers high densities of specific types of voltage gated ion channels and anchors itself to the underlying actin cytoskeleton via βIV Spectrin (Rasband, 2010; Leterrier, 2018). Genes encoding voltage gated sodium channels (VGSCs) that show concentrated distributions within the AIS were also upregulated, as were genes that encode various voltage-gated ion channels expressed within the somatodendritic domain (Lai and Jan, 2006; Hu *et al*., 2009).

Based on the above analysis, we hypothesised that conditional Pax6 deletion affects the development of prethalamic neuronal morphology, the formation of the AIS and the activity of prethalamic neurons. To test these hypotheses, we used the same conditional mutant mouse model as in the RNAseq study and measured the effects in an *in vitro* culture system.

### Pax6 loss caused defects of neurite extension in developing prethalamic neurons

Our protocol for tamoxifen-induced deletion caused Pax6 protein loss from E11.5 onwards in conditional knockouts (Pax6cKOs, CAG^CreER^ Pax6^fl/fl^) (Quintana-Urzainqui *et al*., 2018). No Pax6 protein was detected at the time of dissociation at E13.5 (Figure 1B-B’). Littermates that were heterozygous for the Pax6^fl^ allele were used as the control group (Ctrl, CAG^CreER^ Pax6^fl/+^), as they continue to express Pax6 in a normal pattern (Simpson *et al*., 2009).

Most cultured Ctrl and Pax6cKO prethalamic cells were positive for Tuj1 and neuritogenesis had begun at 1 day in vitro (1DIV) (Figure 2G-G’). Most prethalamic neurons displayed one longest neurite, presumably the developing axon, and several shorter neurites (Figure 2G-L’). At each DIV, we measured: the number of neurites; the length of the longest neurite; the total length of neurites (Figure 2F-F’). Pax6cKO prethalamic neurons had fewer neurites than Ctrl neurons at 1DIV but not after longer culture (Figure 2M). The longest neurites and the total neurite lengths in Pax6cKO prethalamic neurons were shorter than in Ctrls from 1-3 DIV (Figure 2N-O). However, after 3DIV the longest neurites elongated more rapidly in Pax6cKO than in Ctrl neurons and they became significantly longer at 5DIV and 6DIV (Figure 2N). Thus, Pax6cKO prethalamic neurons developed a relatively normal complement of neurites after a delayed start, the longest of which later outstripped their equivalents in Ctrl neurons. Mechanisms limiting axon elongation might be important because most prethalamic neurons project only short distances (Martinez-Ferre and Martinez, 2012; Willis *et al*., 2015), unlike neighbouring thalamic neurons, many of which project long thalamocortical axons (TCAs).

**Figure 2.**
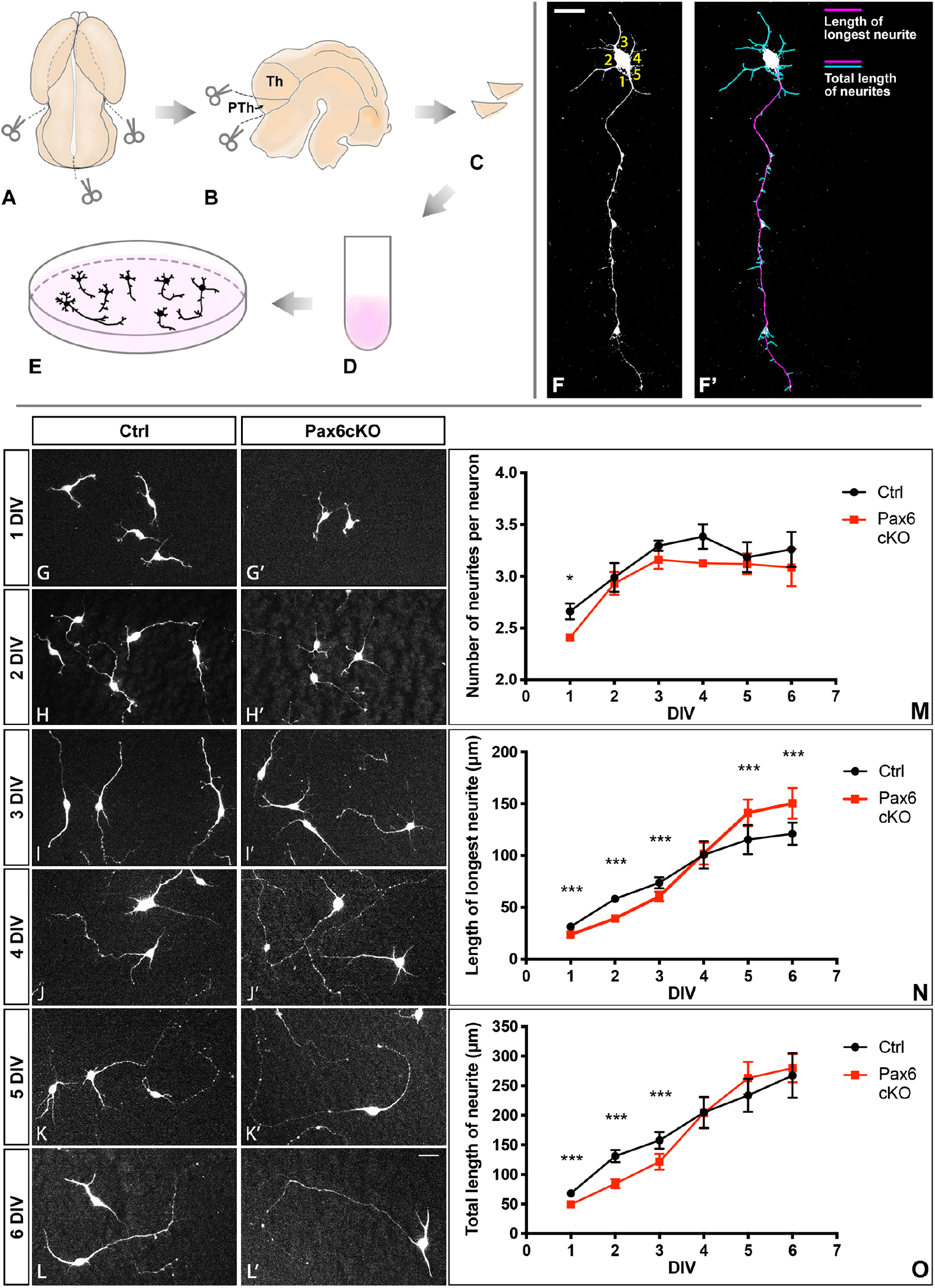
Neuronal morphogenesis of prethalamic neurons in vitro. (A-E) Schematic summary of E13.5 prethalamus dissection for dissociated cell culture. (F-F’) Example of how the number of neurites (F), the length of the longest neurite and the total length of neurites (F’) were measured for each neuron. Scale bar: 20μm (G-L’) Example of prethalamic neuron morphology labelled with Tuj1 from both genotypes on each DIV. Scale bar: 10μm. (M-O) Quantification of the number of neurites (M), length of longest neurite (N) and total length of neurites (O) in Ctrl and Pax6cKO prethalamic neurons cultured 1-6DIV. Mixed-effect model, at least 100 neurons from each genotype were collected from each litter on each of the six DIVs. Three litters in total (n=3). ***: P<0.001, **: 0.001<P<0.01, *: 0.01<P<0.05. Mean with standard error of mean (SEM).

It is unclear what causes delayed neuronal morphogenesis in Pax6cKO prethalamic neurons at early stages of cell culture. Previous studies proposed that the establishment of neuronal polarity can be one of the speed-limiting steps of neuronal morphogenesis in vitro (Bradke and Dotti, 2000; Barnes and Polleux, 2009; Yogev and Shen, 2017). We examined this further, since the RNAseq analysis indicated that Pax6cKO prethalamic cells upregulated expression of genes encoding components of the Par3/6 complex. We used the localised distribution of Par3 as an indicator of neuronal polarity (Nishimura *et al*., 2004). Our results showed no significant differences in the amount and distribution of Par3 within the neurites and the soma between Ctrl and Pax6cKO prethalamic neurons at 3DIV (Supplementary Figure 1). We concluded that the establishment of neuronal polarity was not affected in *Pax6*-deleted prethalamic neurons.

### Pax6 loss altered AIS length and location in developing prethalamic neurons

We investigated AIS formation with immunohistochemistry for AnkG and VGSC in dissociated prethalamic neurons at 7 and 9 DIV.

At both 7DIV and 9DIV, the majority of the prethalamic neurons of both genotypes developed a single AIS (Figure 3A-D, magenta segments) where the profiles of AnkG and VGSC expression (quantified as illustrated in Figure. 3A’-D’) coincided at the proximal end of a neurite (supporting our conclusion that the establishment of neuronal polarity was unaffected in the Pax6cKO prethalamic neurons). We observed a tendency for the AISs to extend further from the soma in Pax6cKO neurons (Figure. 3E-G). The average lengths of the AIS measured with VGSC at 7DIV and with AnkG at 9DIV were significantly higher in Pax6cKO neurons (Figure. 3H, I). In no case were densities of staining (i.e. the average fluorescence intensity per unit length of neurite) with either marker different between the two genotypes (Figure. 3J, K). Figure 3L-M summarises the lengthening and distal extension of the AIS in the prethalamic neurons after 7 and 9 DIV.

**Figure 3.**
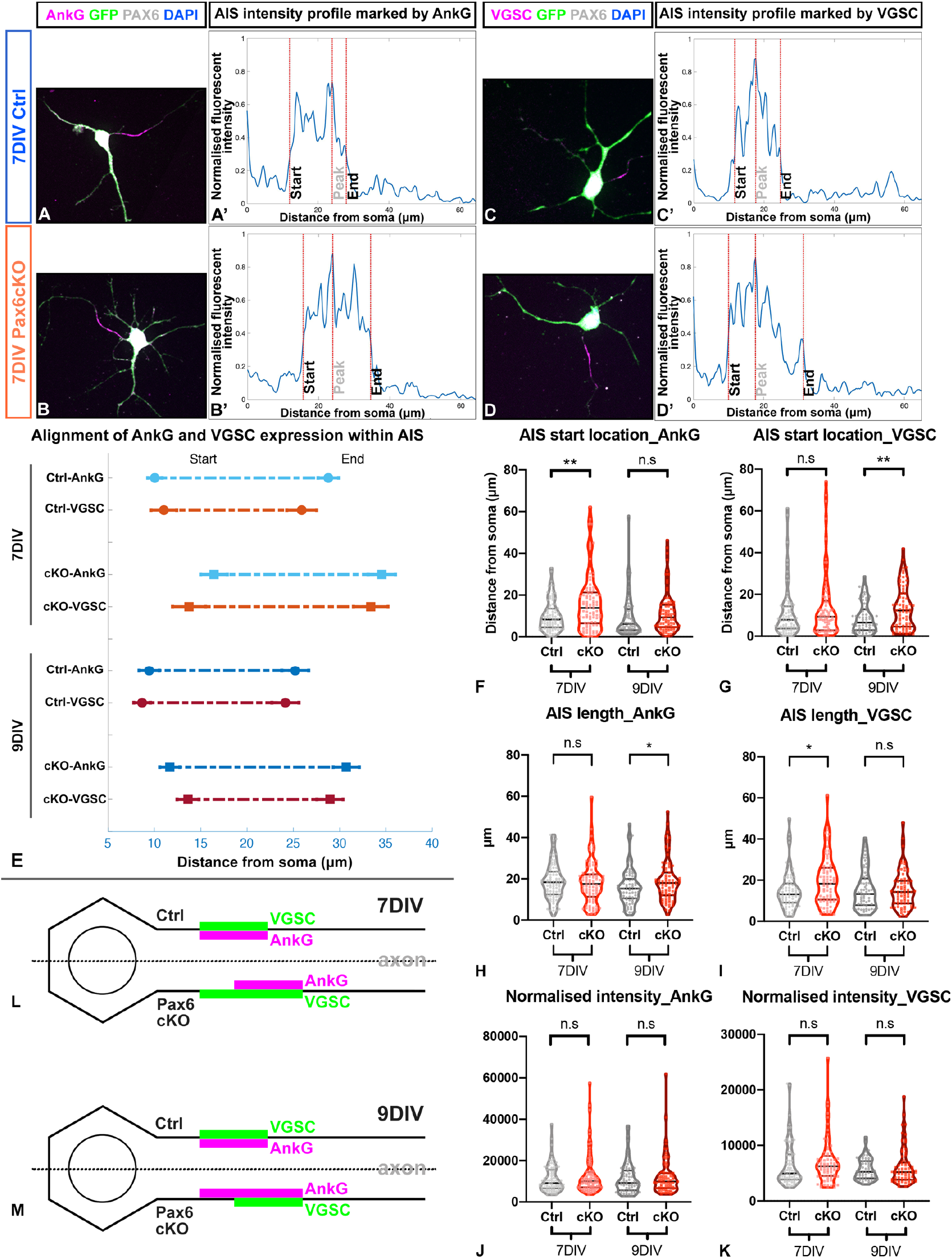
Loss of Pax6 changed the length and location of the AIS in the prethalamic neurons. (A-D) Examples of immunohistochemistry staining showing localised distribution of AnkG or VGSC, marking the AIS, in prethalamic neurons cultured for 7 DIV. Scale bar: 10μm. (A’-D’) Examples of normalised fluorescence intensity of AnkG or VGSC along the axon, termed AIS intensity profile. The peak is defined as where the normalised fluorescence intensity is the highest. The start location (close to the soma) and the end location are defined as where the normalised fluorescence intensities fall to 33% of peak intensity (Grubb and Burrone, 2010). (E) Alignment of the AIS length and location marked by the two markers (AnkG and VGSC) in Ctrl and Pax6cKO prethalamic neurons cultured for 7 and 9 DIV. Comparison of the start location (F-G), length (H-I) and normalised intensity (J-K) of the AIS marked by either AnkG or VGSCs in Ctrl and Pax6cKO prethalamic neurons cultured for 7 and 9 DIV. Length of the AIS is defined as the distance between the start and the end location. (L-M) Schematic summary of changes of AIS length and location in Pax6cKO prethalamic neurons at 7 and 9 DIV. Mixed-effect model, at least 30 neurons from each genotype were collected from each litter on either 7 or 9 DIV. Three litters in total (n=3). ***: P<0.001, **: 0.001<P<0.01, *: 0.01<P<0.05. (E) Mean with SEM. (F-K) Median with interquartile range.

To our knowledge, this is the first evidence that Pax6 is required for the normal formation of AISs. AIS assembly is considered an intrinsic property of neurons, requiring no extracellular or glial-dependent cues (Ogawa and Rasband, 2008). The observed lengthening of the AIS was in line with the upregulated expression of VGSCs and AnkG shown in the RNAseq data, but their distal shift was unexpected. As the master regulator of AIS assembly, AnkG specifies AIS formation and VGSC clustering (Zhou *et al*., 1998; Rasband, 2010), but little is known about the mechanisms that contribute to the enrichment and targeting of AnkG to the proximal axon and specify the location of the AIS (Rasband, 2010; Berger *et al*., 2018; Leterrier, 2018).

### Loss of Pax6 affected the electrophysiological properties of prethalamic neurons

Since the reported changes of the AIS and the expression of voltage-gated ion channels might change neuronal excitability and electrical functions (Grubb and Burrone, 2010; Grubb *et al*., 2011; Kaphzan *et al*., 2011; Höfflin *et al*., 2017; Booker *et al*., 2020), we performed whole-cell patch-clamp on prethalamic neurons cultured for 7 and 9 DIV.

To induce action potentials (APs), we stimulated the prethalamic neurons with small depolarising current steps from -60mV. Prethalamic neurons of both genotypes at both ages were able to fire APs (Figure 4A-B’). From 7 to 9DIV, the resting membrane potentials (RMPs) became significantly more negative in both the Ctrl and Pax6cKO prethalamic neurons, as expected in maturing neurons (Linaro *et al*., 2019), with no significant differences between genotypes (Figure 4C). Figure 4D-E showed how membrane potentials changed in response to specific negative (hyperpolarising) and positive (depolarising) current inputs before rheobases were reached. At 7DIV but not at 9DIV, negative current inputs hyperpolarised membrane potentials significantly more in Pax6cKO than in Ctrl neurons (Figure 4D-E). Although no differences were found at 7DIV (p=0.0549), rheobase became significantly lower in Pax6 cKOs at 9DIV (p=0.0077) (Figure 4F). The AP threshold remained unchanged at both ages (Figure 4G).

**Figure 4.**
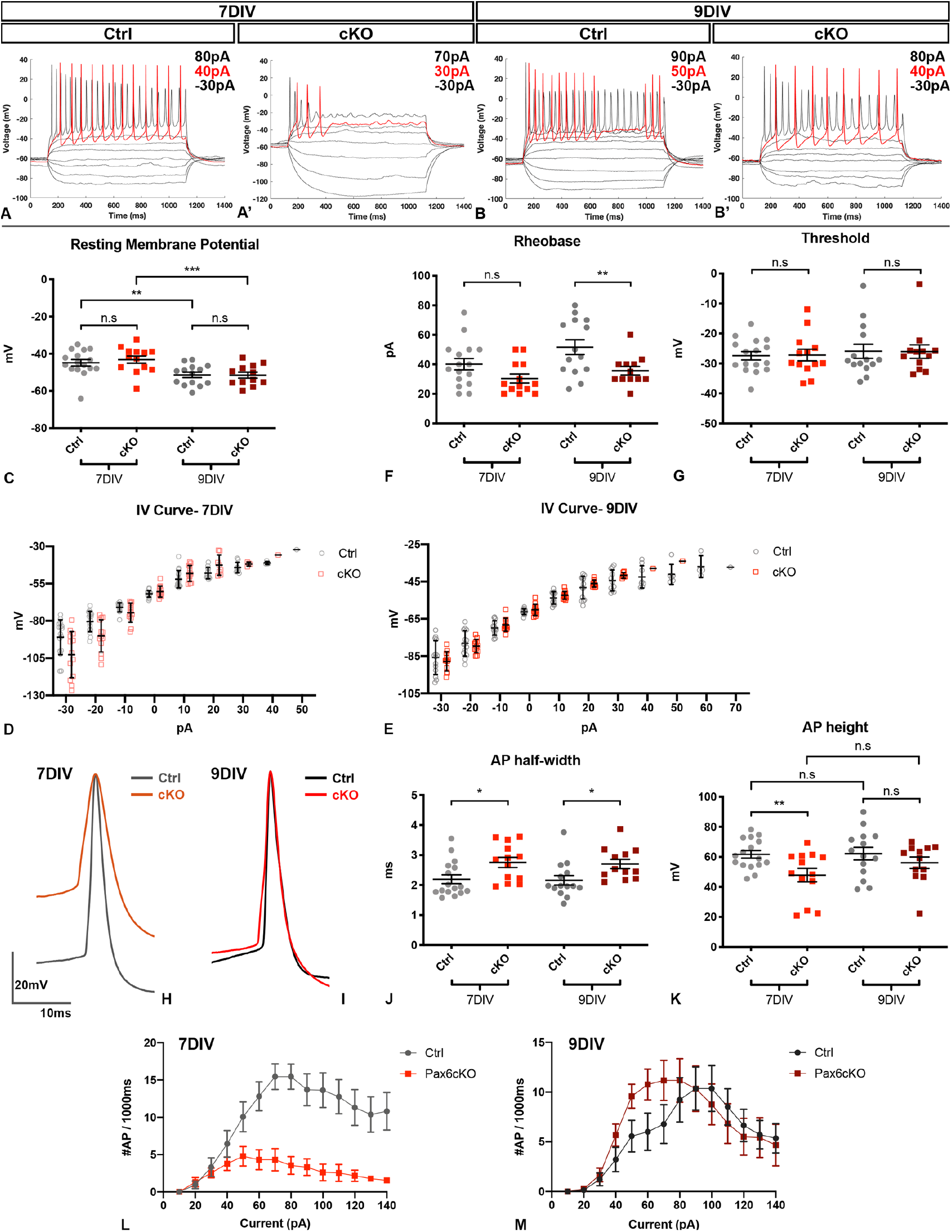
Pax6cKO prethalamic neurons fire APs differently at 7 and 9 DIV. (A-B’) Example of membrane potential changes of Ctrl and Pax6cKO prethalamic neurons responding to depolarising current steps (−30pA to 140pA, 10pA increments, 1000ms). The red traces in each figure showed membrane potential traces at rheobases. (C, F, G) Comparison of resting membrane potential, rheobase and threshold potential in Ctrl and Pax6cKO prethalamic neurons cultured for 7 and 9 DIV. (D-E) I-V curves in Ctrl and Pax6cKO prethalamic neurons cultured for 7 and 9 DIV. (H-I) Aligned and overlayed traces of the first APs fired at rheobases by the Ctrl and Pax6cKO prethalamic neurons cultured for 7 and 9 DIV. (J-K) Comparison of the half-width and height of the first AP at rheobases in Ctrl and Pax6cKO prethalamic neurons cultured for 7 and 9 DIV. (L-M) Current frequency plot for Ctrl and Pax6cKO prethalamic neurons cultured for 7 and 9DIV, indicating differences in AP firing frequency in response to applied current steps. Mixed effect model, 16 neurons from Ctrl 7DIV, 13 neurons from Pax6cKO 7DIV, 14 neurons from Ctrl 9DIV, 12 neurons from Pax6cKO 9DIV. 4 litters for each genotype at both DIVs (n=4). ***: P<0.001, **: 0.001<P<0.01, *: 0.01<P<0.05. (C-G, J-K) Mean with SEM. (D-E) Mean with SD.

We examined the waveforms of the first APs fired by the Ctrl and the Pax6cKO prethalamic neurons in response to the current stimulus at their rheobases. Figure 4H-I shows the overlayed example traces of the first APs shown in Figure 4A-B’. AP halfwidth was significantly broader in Pax6cKOs at both ages and AP height was significantly lower in Pax6cKOs at 7DIV (Figure 4J-K).

Prethalamic neurons from both genotypes were able to fire multiple APs after rheobases were reached but, at 7DIV, Pax6cKO prethalamic neurons showed a significant reduction in maximum AP number (Figure 4L). The highest AP firing frequency in the Pax6cKO neurons at 7DIV (7± 1.4Hz) was significantly lower than that in the Ctrl neurons (18.6 ± 1.9Hz). This phenotype was recovered at 9DIV, when Pax6cKO and Ctrl neurons fired comparable maximum numbers of APs (Figure 4M). At 9DIV, Pax6 loss also caused a leftward shift in the current-frequency response (Figure 4M), indicating that a similar number of APs were generated using a lower amplitude of current stimulus in the Pax6cKO neurons.

Therefore, Pax6 loss can affect the activity of the prethalamic neurons, resulting in changes in their excitability level, the waveforms of somatic APs and their ability to fire APs repetitively.

The distal shift of the AIS in Pax6cKO neurons might have contributed to the widening of the AP waveforms recorded from their soma as these recordings would have been mainly the backpropagated APs generated at the AIS (Kole, Letzkus and Stuart, 2007) and increasing the distance between the AIS and the soma would increase voltage attenuation, resulting in wider and slower APs. The underlying causes for the decrease in rheobase in the Pax6cKO prethalamic neurons are less clear. Changes in the AIS length and location could be a factor since previous studies have shown that both increased length and distal relocation of the AIS can promote excitability (Grubb and Burrone, 2010; Buffington and Rasband, 2011; Kaphzan *et al*., 2011; Höfflin *et al*., 2017; Jamann, Jordan and Engelhardt, 2018; Goethals and Brette, 2020). Changes of AIS geometry alone are unlikely to provide a full explanation, however, since theoretical modelling suggests that they would produce a modest decrease (a few mV) in threshold potential (Goethals and Brette, 2020), for which we found no evidence. Therefore, it is likely that changes outside the AIS contributed to the decrease in rheobase in Pax6cKO prethalamic neurons. The surprising observation that 7DIV Pax6cKO prethalamic neurons fired significantly fewer APs and entered the state of depolarization block with significantly lower amplitude of current stimulus despite having significantly longer AISs might be explained in two ways. First, there might be defects at the AISs; second, RNAseq data suggested upregulation of several voltage-gated potassium channels belonging to the Kv4 family (Kv4.1(KCND1), Kv4.2(KCND2) and Kv4.3(KCND3), Supplementary Table 2), which can efficiently reduce repetitive firing (Fransén and Tigerholm, 2010; Hermanstyne *et al*., 2017; Kim *et al*., 2020).

In conclusion, our results suggest that the consequences of Pax6 loss from the prethalamic neuroepithelium include the generation of neurons with an abnormal developmental potential. Mutant prethalamic neurons show abnormal axonal extension, AISs and somatic AP waveforms and excitability, which might impact on prethalamus-thalamus circuit formation and the functional development of the nervous system.

## MATERIALS AND METHODS

### Experimental Model and Subject Details

#### Mice colony maintenance and transgenic lines

Mouse lines used to generate tamoxifen-induced deletion of Pax6 throughout the embryo were as described in (Quintana-Urzainqui *et al*., 2018). Pregnant mice were given 10mg of tamoxifen (Sigma) by oral gavage on embryonic day 9.5 (E9.5) to induce Pax6loxP deletion and embryos were collected on E13.5. Embryos heterozygous for the Pax6flox allele (Pax6^fl/+^; CAGG^CreER^) were used as controls (Ctrl) as previous studies have shown no detectable defects in the forebrain of Pax6^fl/+^ embryos (Simpson *et al*., 2009). Embryos carrying two copies of the floxed Pax6 allele (Pax6^fl/fl^; CAGG^CreER^) were the experimental conditional knock-out (cKO) group.

For staging of embryos, the first day the vaginal plug was detected was considered as embryonic day 0.5 (E0.5).

All animal husbandry was conducted in accordance with the UK Animal (Scientific Procedures) Act 1986 regulations and all procedures were approved by Edinburgh University’s Animal Ethics Committee.

### Method Details

#### Dissociated prethalamic cell culture preparation

##### Dissection of the prethalamus

To dissect the prethalamus, embryos from E13.5 were collected and decapitated. The neural tube was separated from the epidermal and mesodermal tissue and cut in half along the dorsal and ventral midline (Figure 2A-B). From E9.5, morphologic segmentation of the diencephalon starts and the diencephalic prosomeres become apparent from E10-11 as ventricular ridges and lateral wall bulges appear (18). These morphological landmarks are used to distinguish prethalamus from the surrounding tissue (the thalamus and the eminentia thalami) during dissection (Figure 2C).

##### Dissociated cell culture

After being cut out, the prethalamus from the two halves of the neural tube of the same embryo were put together and chopped into smaller pieces for dissociated cell culture using the Papain Dissociation System (Worthington Biochemical Corp.) according to the manufacturer’s protocol (Figure 2C-E). The prethalamus from each embryo was dissociated and cultured individually. The number of cells obtained after dissociation was measured using a haemocytometer. Additionally, trypan blue staining was used to determine the ratio of viable to damaged cells. To adjust the plating density of the cell culture, cells obtained after dissociation were resuspended using a certain amount of culture medium (Advanced DMEM/Neurobasal medium 1:1, supplemented with N2 (100x) and B27 (50x) neural supplement, Thermo Fisher Scientific). 130μl of the culture medium containing the desired number of resuspended cells was then added onto the 9mm circular coverslips (Fischer Scientific) coated with Poly-L-lysine and Laminin (Thermo Fisher Scientific). Due to the surface tension of the culture medium, the culture medium containing the dissociated cells would stay within and fill up the realm of the coverslips. The cell cultures were then incubated at 37°C with 5% CO2 for 1 hour to allow the cells to attach to the coverslips. In this way, all the cells obtained after the dissociation was retained within the coverslip, and the exact cell density of plating could be calculated using the total amount of cells divided by the surface areas of the coverslips being used. The plating density of prethalamic cell culture used for studying neuronal morphogenesis and AIS formation was 20 cells/mm^2^, and the plating density of prethalamic cell culture used for electrophysiological recordings was 600 cells/mm^2^. After 1hr, 240μl of the culture medium was then added into each well of the 48-well plate (Greiner Bio-One) that contains the coverslip. The dissociated prethalamic cells were cultured for 1-9 days. Light microscopy was used to monitor the condition of cell cultures daily.

#### Genotyping

We dissected tissue from the tails of each embryo, extracted DNA and performed PCR amplification to detect the alleles of interest.

For the detection of the floxed Pax6 allele, PCR reaction was performed in a final volume of 25μl containing 1.5μl of extracted DNA, 0.5mM primer mix (Simpson et al. 2009, forward primer: 5’-AAA TGG GGG TGA AGT GTG AG-3’; reverse primer: 5’-TGC ATG TTG CCT GAA AGA AG-3’), 0.5 mM dNTPs mix, 1X PCR reaction buffer and 5U/μl Taq DNA Polymerase (Qiagen). PCR was performed with 35 cycles and a Tm of 59°C. The PCR product was subsequently run in a 2% agarose gel. Wild type allele results in a fragment of 156bp and floxed allele fragment was 195bp, therefore two bands indicated the heterozygous condition (used as controls) and one strong 195bp band identified the homozygous floxed allele condition (Pax6cKOs).

#### Histological processing and imaging

##### Sample processing for immunohistochemistry

Cryo-sections were obtained following methods described in (Quintana-Urzainqui *et al*., 2018).

Cell culture samples were obtained by removing the coverslips containing the prethalamic neurons from the culture medium. 1x phosphate buffered saline (PBS) warmed to 37°C were used to rinse the prethalamic neurons for 3 times. For cell culture samples used in neuronal morphology and polarity studies, the prethalamic neurons were fixed in 4% PFA for 20 minutes. For cell culture samples used in AIS study, the prethalamic neurons were fixed in 2% PFA/4% sucrose for 10 minutes to prevent degradation of AnkG protein by the fixative. The cell culture samples were further rinsed with 1x PBS and kept in 1x PBS at 4°C until processed.

##### Fluorescent immunohistochemistry

Fluorescent immunohistochemistry on cryo-sections were performed as described in (Quintana-Urzainqui *et al*., 2018).

Fluorescent immunohistochemistry on cell culture were performed as follows: fixed prethalamic neurons were rinsed with 0.1% Triton X-100 in 1xPBS (0.1% PBST) and permeabilised in 0.5% Triton X-100 in 1x PBS (0.5% PBST) for 10 minutes. Then cells were washed with 0.1% PBST for three times and further blocked with blocking solution (20% goat or donkey serum in 0.1% PBST) for 2 hours. Then the blocking serum containing primary antibodies was added to the cells for overnight incubation. On the second day, cells were washed with 0.1% PBST and further incubated with blocking serum containing the corresponding secondary antibodies (Streptavidin Alexa Fluor™ 488, 546 or 647 conjugates; Thermo Fisher Scientific) for 1 hour. Cells were then washed with 1xPBS, and further incubated with DAPI (Thermo Fisher Scientific) for counterstaining of the nucleus. The coverslips were mounted in ProLong Gold Antifade Mountant (Thermo Fisher Scientific) for further imaging.

##### Imaging

Fluorescence images of the dissociated cells in the neuronal morphogenesis study were taken using a Leica DM5500B automated upright microscope connected to a DFC360FX camera. The route of acquiring images started from the upper-left corner towards the lower-right of the 9mm circular coverslip. Every image being taken was from an adjacent visual field of the previous image to cover as many cells on the coverslip as possible and without imaging the same cells twice.

Fluorescence images of the dissociated cells in the PAR3 study were taken using the Zeiss LSM800 confocal microscope with Airy Scan.

Fluorescence images of the dissociated cells in the AIS study were taken using the Andor Revolution XDi Spinning disk confocal microscope.

#### Whole-cell patch-clamp recording

For electrophysiological recordings on the dissociated prethalamic neurons cultured for 7 and 9 DIV, coverslips with attached prethalamic neurons were transferred to a submerged recording chamber perfused with carbonated ACSF (in mM: 150 NaCl, 2.8 KCl, 10 HEPES, 2 CaCl2, 1 MgCl2, 10 Glucose, pH7.3), at a flow rate of 4-6ml/min at 22-23°C. Prethalamic neurons were visualised with a digital camera (SciCam Pro, Scientifica) mounted on an upright microscope (BX61-WI, Olympus) and a 40x water-immersion objective lens (1.0 N.A., Olympus). Whole-cell patch-clamp recordings were performed with a Multiclamp 700B amplifier (Molecular Devices), filtered at 10KHz with the built-in 4-pole Bessel Filter and digitised using a Digidata 1440A digitiser board (Molecular Devices) at 20kHz. Recording pipettes were pulled from borosilicate glass capillaries (Havard Apparatus, 30-0060) on a horizontal electrode puller (P-97, Sutter Instruments) to a tip resistance of 4-6 MΩ. Recording pipettes were filled with K-gluconate-based internal solution (in mM: 130 K-gluconate, 4 Glucose, 10 HEPES, 0.1 EGTA, 0.025 CaCl2, 20 Sucrose, pH=7.2, 290-300mOsm). Recording pipettes were positioned with a micromanipulator (Scientifica PatchStar). Data were acquired from cells with access resistance < 25MΩ. pClamp 10 (Axon Instruments) was used to generate the various analogue waveforms to control the amplifier and record the traces.

Resting membrane potential of the prethalamic neurons was recorded with current clamped with no current input (I=0 pA) for 30 seconds. Constant current was injected to hold the cells close to -60mV and I-V curve, rheobase and threshold was assessed by current injections from -30 to +140 pA for 1000 ms (10 pA steps). All AP properties were determined from the first AP elicited at rheobase. Analysis of electrophysiological data was performed offline using a custom-written Matlab script kindly provided by Dr Adam Jackson, blind to genotype.

#### Image analysis

##### Measurement of neurite length

Measurement of neurite length was performed using the freehand line tool in the FIJI package of ImageJ. Whole morphology of the neurons was marked by fluorescent immunohistochemistry reacting with the Tuj1 antibody, which labels the neuron-specific Class III β-tubulin of the cytoskeleton. For each Tuj1-positive cell, three parameters - the number of neurites, the length of the longest neurite and the total length of neurites were measured. A neurite was defined as a stable protrusion from the soma with strong Tuj1 staining. Protrusions from the soma that were thin and had faint Tuj1 staining were considered as either artefacts or filopodia, which mainly consist of F-actin and were not considered as neurites. The length of a neurite was measured by tracing down the neurite from the edge of the soma to the most distal edge of Tuj1 staining. Such measurement was performed for every neurite of each neuron analysed. The length of the longest neurite was the highest reading among these measurements of each neuron. All the protrusions, positive for Tuj1 staining that stemmed from the neurites were considered as branches. The length of each branch was measured from where it stemmed from the neurite to its furthest edge of Tuj1 staining. The total length of neurites for each neuron was calculated as the sum of the length of all the neurites and branches.

##### Measurement of PAR3 distribution

Measurement was performed with the IMARIS software (Bitplane, version 9.1.2). Dissociated prethalamic neurons cultured for 3DIV were marked by fluorescent immunohistochemistry reacting with antibodies for PAR3, GFP for whole-cell morphology and DAPI for counterstaining of the nucleus.

The surface function of IMARIS was utilised to create a new GFP channel. To do so, the signal intensity threshold was adjusted manually based on the specific situation of each neuron so that IMARIS was able to detect the space where the GFP signal is above that set threshold. This threshold was determined by fitting and adjusting different threshold values so that the GFP-positive space IMARIS detected optimally matched the actual cytoplasmic volume and included almost all the neurites and cytoplasmic protrusions. Places where their GFP signals were above this threshold but resided outside of the cytoplasm were manually deleted. In this way, a new GFP channel was created, which only included and highly resembled the entire cytoplasm of the neurons. Then a new PAR3 channel was created using the new GFP channel as the template of the cytoplasmic volume to exclude any Par3 staining outside of the cytoplasm. With this new PAR3 channel, we could then detect the highest intensity of PAR3 expression within the cytoplasm. Different thresholds were set for IMARIS to detect the cytoplasmic volumes in which the PAR3 intensities were above 75%, 50%, 25% and 10% of its own highest intensity value. This allowed us to quantify the cytoplasmic volume that had higher expression levels of PAR3 at each of these thresholds, and also visualise the distribution of different intensities of PAR3 within the cytoplasm.

##### Measurement of the intensity profile of AIS

The intensity profile of AIS was generated by tracing down the neurite bearing the localised expression of either AnkG or VGSC, 80μm from the edge of the soma, using the freehand line tool in the FIJI package of ImageJ, resulting in a graph with the level of AnkG or VGSC expression at each pixel from the soma corresponding to the specific distance from the soma.

### Quantification and Statistical Analysis

All experiments were performed blind to genotype. All data was analysed with either a linear mixed-effect model (LMM) or its generalised form (GLMM), unless stated otherwise. The variability due to random effects (animal, litter) was taken into account, allowing for the calculation of the genotype effect size. Where reported, statistical significance was assumed if p<0.05.

### Key Resource Table

| Reagent or Resource | Source | Identifier |
| --- | --- | --- |
| **Antibodies** |  |  |
| Rabbit polyclonal anti-Pax6 (1:200) | BioLegend | Cat#901301, RRID: AB_2565003 |
| Mouse monoclonal anti-Pax6 (1:10) | A gift from Prof. V van Heyningen, AD2.38<br>Simpson *et al.*, 2009 | N/A |
| Mouse monoclonal anti-Beta III tubulin (Tuj1) (1:200) | Abcam | Cat# ab18207, RRID: AB_444319 |
| Mouse monoclonal anti-Ankyrin-G, clone N106/36 (1:200) | Antibodies Inc. | Cat# 75-146 |
| Monoclonal Anti-Sodium Channel, Pan antibody produced in mouse (1:100) | Sigma-Aldrich | Cat# S8809-.1MG |
| Rabbit polyclonal Anti-Partitioning-defective 3 Antibody (1:100) | Millipore | Cat# 07-330 |
| Goat polyclonal Anti-GFP antibody (1:200) | Abcam | Cat# ab6673 |
| Chicken polyclonal Anti-GFP antibody (1:200) | Abcam | Cat# ab13970 |
| Goat anti-mouse biotinylated secondary antibody (1:200) | Vector laboratories | Cat# BA-9200, RRID:AB_2336171 |
| Goat anti-rabbit biotinylated secondary antibody (1:200) | Vector laboratories | Cat# BA-1000, RRID:AB_2313606 |
| Donkey anti-mouse Alexa 568 secondary antibody (1:200) | Thermo Fisher Scientific | Cat# A10037, RRID:AB_2534013 |
| Donkey anti-rabbit Alexa 647 secondary antibody (1:200) | Invitrogen | Cat # A-31573 |
| Donkey anti-chicken Alexa 488 secondary antibody (1:200) | Strattech Scientific Limited | Cat# 703-545-155-JIR |
| Donkey anti-rabbit Alexa 568 secondary antibody (1:200) | Invitrogen | Cat # A10042 |
| Donkey anti-goat Alexa 488 secondary antibody (1:200) | Invitrogen | Cat # A-11055 |
| Donkey anti-rat Alexa 488 secondary antibody (1:200) | Thermo Fisher Scientific | Cat# A-21208, RRID:AB_141709 |
| Streptavidin, Alexa Fluor 488 conjugate antibody (1:200) | Thermo Fisher Scientific | Cat# S11223, RRID:AB_2336881 |
| **Chemicals, Peptides, and Recombinant Proteins** |  |  |
| Tamoxifen | Sigma-Aldrich | T5648; CAS: 10540-29-1 |
| **Critical Commercial Assays** |  |  |
| DAB peroxidase substrate kit | Vector Laboratories | SK4100 |
| Papain Dissociation System | Worthington Biochemical Corp | Cat# LK003150 |
| **Experimental Models:**<br>**Organisms/Strains** |  |  |
| Mouse: CAGGCre-ERTM; RCE (GFP) | This paper Hayashi and McMahon, 2002 Sousa et al., 2009 | N/A |
| Mouse: Pax6loxP | Simpson et al., 2009 | N/A |
| **Software and Algorithms** |  |  |
| Graphpad Prism 8 | Graphpad software |  |
| pClamp | Molecular Devices |  |
| Matlab_R2018b | Mathworks |  |
| Fiji (Image J) | Schindelin et al., 2012 | https://fiji.sc/ |
| IMARIS 9.1.2 | Bitplane |  |

## Acknowledgement

The authors would like to thank Dr Adam Jackson and Dr Javier Moral-Sanz for assistance with experimental set up for whole-cell patch-clamp recordings, Dr Michael Daw and Dr Sam Booker for comments on the manuscript and all of the members of the Developmental Biology User Group for helpful discussion.

## Author Contributions

T.T., I.Q.-U., T.P., and D.J.P. were involved in overall conceptualization, data interpretation, and manuscript preparation. T.T. carried out wet-laboratory experiments; Z.K. and T.T. did the bioinformatics and statistics. D.J.P. acquired funding and I.Q.-U., T.P. and D.J.P. supervised the work.

## Resource Availability

### Lead contact

Further information and requests for resources and reagents should be directed to and will be fulfilled by the Lead Contact

### Materials Availability

This study did not generate new unique reagents.

### Data and Code Availability

All data and code generated in this study will be made available upon reasonable request.

The published article (Quintana-Urzainqui *et al*., 2018) includes all the RNAseq dataset analysed during this study.

## Supplementary Information

**Supplemental Table 1.** List of significantly up- and downregulated (adjusted p<0.05) genes and GO terms in the E13.5 prethalamus after induced acute Pax6 deletion at E9.5. Related to Figure 1.

| Genes | LFC | P value | Description |
| --- | --- | --- | --- |
| **GEFs for Rac** |  |  |  |
| Dock4 | 1.03 | 3.39E-10 | dedicator of cytokinesis 4 |
| Dock2 | 0.86 | 0.00196 | dedicator of cytokinesis 2 |
| Dock3 | 0.68 | 0.00300 | dedicator of cytokinesis 3 |
| Dock9 | 0.66 | 0.00671 | dedicator of cytokinesis 9 |
| Dock10 | 0.76 | 0.00339 | dedicator of cytokinesis 10 |
| Dock8 | 0.95 | 0.00178 | dedicator of cytokinesis 8 |
| **Scaffold for GEFs** |  |  |  |
| Elmo1 | 0.87 | 2.21E-10 | engulfment and cell motility 1 |
| **Scaffold for CDC42** |  |  |  |
| Fgd1 | 0.57 | 0.00362 | FYVE, RhoGEF and PH domain containing 1 |
| Fgd5 | 1.04 | 3.11E-05 | FYVE, RhoGEF and PH domain containing 5 |
| Fgd2 | 0.67 | 0.00234 | FYVE, RhoGEF and PH domain containing 2 |
| Fgd6 | 0.65 | 0.00503 | FYVE, RhoGEF and PH domain containing 6 |
| **GAP for Rho and Rac** |  |  |  |
| Srgap3 | 0.92 | 0.00086 | SLIT-ROBO Rho GTPase activating protein 3 |
| **Rac** |  |  |  |
| Rnd1 | 0.77 | 2.55E-07 | Rho family GTPase 1 |
| **Actin-bundling** |  |  |  |
| Ablim1 | 0.59 | 0.00253 | actin-binding LIM protein 1 |
| Ablim3 | 1.19 | 7.20E-07 | actin binding LIM protein family, member 3 |
| Ablim2 | 0.91 | 0.00342 | actin-binding LIM protein 2 |
| Strip2 | 1.80 | 4.32E-07 | striatin interacting protein 2 |
| **Anti-capping for actin** |  |  |  |
| Enah | 0.67 | 0.00391 | enabled homolog (Drosophila) |
| **Membrane curvature** |  |  |  |
| Pacsin1 | 0.81 | 0.00546 | protein kinase C and casein kinase substrate in neurons 1 |
| **Microtubule polymerisation/bundling** |  |  |  |
| Mapt | 1.16 | 4.90E-08 | microtubule-associated protein tau |
| Map6 | 0.70 | 0.00732 | microtubule-associated protein 6 |
| Map9 | 0.60 | 0.00746 | microtubule-associated protein 9 |
| Map2 | 0.32 | 0.01108 | microtubule-associated protein 2 |
| Mark1 | 0.78 | 0.00014 | MAP/microtubule affinity-regulating kinase 1 |
| Mark4 | 0.76 | 0.00720 | MAP/microtubule affinity-regulating kinase 4 |
| Apc2 | 0.86 | 0.00485 | adenomatosis polyposis coli 2 |
| Apc | 0.78 | 0.00820 | adenomatosis polyposis coli |
| Crmp1 | 0.32 | 0.00312 | collapsin response mediator protein 1 |
| Dpysl5 | 0.66 | 0.00500 | dihydropyrimidinase-like 5 |
| **Par3/6 complex component** |  |  |  |
| Prkcz | 0.37 | 0.00431 | protein kinase C, zeta |
| Kif3a | 0.45 | 0.00484 | kinesin family member 3A |
| Tiam1 | 0.66 | 0.00821 | T cell lymphoma invasion and metastasis 1 |
| Arhgap35 | 0.59 | 0.01060 | Rho GTPase activating protein 35 |
| Smurf1 | 0.53 | 0.00328 | SMAD specific E3 ubiquitin protein ligase 1 |
| Pard3b | 0.80 | 0.00822 | par-3 family cell polarity regulator beta |
| Apc | 0.78 | 0.00820 | adenomatosis polyposis coli |
| **Cytoskeleton protein in the axon initial segment** |  |  |  |
| Ank3 | 1.19 | 0.00105 | ankyrin 3, epithelial |
| Sptb | 1.17 | 0.00054 | spectrin beta, erythrocytic |
| Sptbn1 | 1.03 | 0.00096 | spectrin beta, non-erythrocytic 1 |
| Sptbn4 | 1.36 | 4.52E-06 | spectrin beta, non-erythrocytic 4 |
| **Voltage-gated Sodium channels (VGSCs)** |  |  |  |
| Scn1a | 0.77 | 0.00123 | sodium channel, voltage-gated, type I, alpha |
| Scn3b | 0.72 | 2.96E-06 | sodium channel, voltage-gated, type III, beta |
| Scn2b | 0.58 | 0.00118 | sodium channel, voltage-gated, type II, beta |
| Scn3a | 0.39 | 0.00502 | sodium channel, voltage-gated, type III, alpha |
| **Rapidly inactivating A-type K<sup>+</sup> channel** |  |  |  |
| Kcna4 | 1.87 | 1.40E-16 | potassium voltage-gated channel, shaker-related subfamily, member 4 |
| Kcnc3 | 1.02 | 0.00040 | potassium voltage gated channel, Shaw-related subfamily, member 3 |
| Kcnc4 | 0.79 | 0.00252 | potassium voltage gated channel, Shaw-related subfamily, member 4 |
| Kcnd1 | 0.61 | 0.00769 | potassium voltage-gated channel, Shal-related family, member 1 |
| Kcnd3 | 1.02 | 3.78E-10 | potassium voltage-gated channel, Shal-related family, member 3 |
| Kcnd2 | 0.84 | 9.81E-07 | potassium voltage-gated channel, Shal-related family, member 2 |
| **Voltage-gated Ca<sup>2+</sup> channels** |  |  |  |
| Cacna1e | 1.09 | 0.00012 | calcium channel, voltage-dependent, R type, alpha 1E subunit |
| Cacna2d2 | 0.68 | 0.00930 | calcium channel, voltage-dependent, alpha 2/delta subunit 2 |
| Cacna1a | 0.61 | 0.00114 | calcium channel, voltage-dependent, P/Q type, alpha 1A subunit |
| Cacna2d4 | 1.87 | 2.98E-08 | calcium channel, voltage-dependent, alpha 2/delta subunit 4 |
| Cacna1g | 1.21 | 4.11E-07 | calcium channel, voltage-dependent, T type, alpha 1G subunit |
| **Hyperpolarization-activated cyclic nucleotide-gated (HCN) channels** |  |  |  |
| Hcn2 | 0.95 | 1.12E-05 | hyperpolarization-activated, cyclic nucleotide-gated K<sup>+</sup> 2 |
| Hcn4 | 0.70 | 0.00481 | hyperpolarization-activated, cyclic nucleotide-gated K<sup>+</sup> 4 |
| Hcn3 | 0.40 | 0.00852 | hyperpolarization-activated, cyclic nucleotide-gated K<sup>+</sup> 3 |

**Supplementary Table 2.** List of genes found in the neuronal morphogenesis and ion transport GO terms that encode proteins with specific neuronal functions. LFC, log-fold change.

**Supplementary Figure 1.**
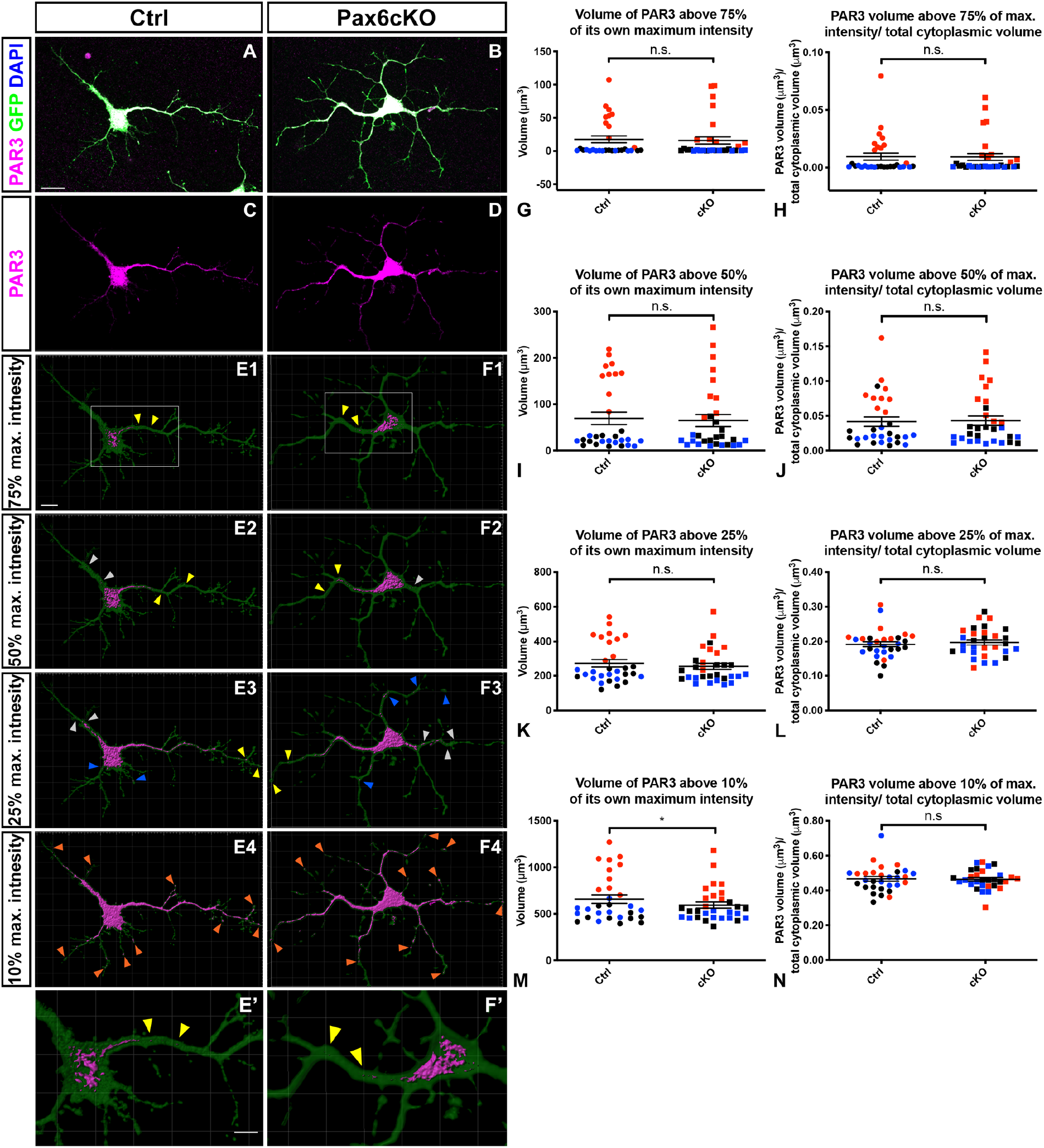
Distribution of Par3 proteins within the developing prethalamic neurons at 3DIV. **(A, B)** Prethalamic neurons from the Ctrl (A) and Pax6cKO (B) embryos were cultured for 3DIV and stained for Par3, GFP and DAPI. **(C, D)** Par3 expression within the cytoplasm of the neurons. **(E1-F1)** Par3 expression within the cytoplasm when the intensity threshold was set at 75% of the maximum intensity of the Par3 channel. The majority of the Par3 volumes were distributed within the soma. However, Par3 volumes were also found in the stem of the longest neurites (yellow arrowheads). **(E2-F2)** Par3 expression within the cytoplasm when the intensity threshold was set at 50% of the maximum intensity of the Par3 channel. More Par3 volumes were detected in the soma, the stem of the longest neurites (yellow arrowheads), as well as in other neurites (grey arrowheads). **(E3-F3)** Par3 expression within the cytoplasm when the intensity threshold was set at 25% of the maximum intensity of the Par3 channel. Par3 volumes within the cytoplasm continued to increase. Par3 volumes could be found at the tip of the longest neurites (yellow arrowheads) and other neurites (grey and blue arrowheads). **(E4-F4)** Par3 expression within the cytoplasm when the intensity threshold was set at 10% of the maximum intensity of the Par3 channel. PAR3 volumes had filled up the cytoplasm and could be found at almost all the tip of the neurites and protrusions (orange arrowheads). (E’-F’) Zoom in on the boxed area in E1 and F1. Scale bars: A-D, 15μm, E1-F5, 10μm, E’-F’, 5μm. (G-N) Comparison of Par3 volumes and percentage of Par3 volumes against the cytoplasmic volume between the Ctrl and the Pax6cKO prethalamic neurons. 10 neurons from the Ctrl and Pax6cKO genotypes were measured. Three litters in total (n=3). Each dot (data point) represented the value of Par3 volume (G, I, K, M) or Par3 volume/ total cytoplasmic volume (H, J, L, N) from each neuron at a specific threshold. The data points of the neurons from the same litter were marked with the same colour. Data were analysed with mixed-effect model. Mean ± SEM. *: P<0.05.

